# Scalable parametric encoding of multiple modalities

**DOI:** 10.1101/2021.07.09.451779

**Authors:** David Banh, Alan Huang

## Abstract

A flexible model is introduced which shares ideas with the Autoencoder, Canonical Correlation Analysis, Singular Value Decomposition, and Procrustes Analysis. It is proposed to find relevant maps to transform multiple datasets of various types from one modality to another. Here, the Generative Encoder is used to transform spatial gene expression from breast tissue, to the images of histology tissue measured with Spatial Transcriptomics. The model is directly interpretable given all parameters are linked to the data space. It is scalable on Big Data, training reasonably on several thousand RGB images of 100 by 100 pixels in under an hour, which equates to 30,000 pixel features per sample image.

## 1 Introduction

Data transform functions aim to map a set of points from one space to another space more useful for further analysis. Transformations are of immense utility, from the kernel trick in Support Vector Machines using a Gaussian transformation [1], to the non-linear composition of functions in neural networks. [2].

Ideas from linear algebra can also have functions acting as transformations. One involving Matrix Projection [3]: used for projecting one matrix onto another second vector or matrix such as in regression, and another being the traditional Procrustes Analysis [4]: used for matching shapes represented by point clouds via matrix rotations and scaling.

State of the art methods that evaluate both the sample and feature spaces appear recently. In 2021, Zhou and Troyanskaya formulate the GraphDR method which projects to a latent space via linearly interpretable parameters that operate on both the samples and features [5]. In 2019, Tarashansky et al with Self-Assembling Manifolds (SAM) use a nearest-neighbour projection method that operates on the samples and features to generate a manifold on the points for further dimensionality reduction and visualisation. [6].

These methods show that the sample space holds vital information that describes nearest-neighbours in a local space, with recent developments also showing the importance of mutual data neighbourhoods [7]. Combining both the data space of samples and features using the ideas of Mutual Nearest-Neighbours with Canonical Correlation Analysis respectively, has led to recent developments in multi-modal integration in single cell analysis [8].

The method developed and proposed in this investigation is similar to the quasi-linear method GraphDR, where model design is chosen such that parameters operate on both the sample and feature spaces. The parameters learn functions which encode both the sample and feature spaces of any dataset, fitting any multiple number of datasets into a shared encoded subspace. From only the learned parameters, the dataset can be decoded to be fully expressed as a generative model. The mapping from the dataset to a lower dimensional space via encoding and mapped back to the original dataset via decoding gives it the style of a Generative Encoder.

## 2 Method

The aim of Generative Encoding is to integrate data from several datasets, *Y, X* and *A*, into an aligned and reduced dimensional space *d* via a transformation. An example of Generative Encoding that is implemented here:

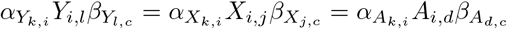

Here, *k* and *c* are the dimensions of the reduced dimensional encoding. The sample size *i* is shared between the datasets *Y, X*, and *A*. However, each dataset contains differing feature sizes *l, j, d*. For example, this could be of a single cell experiment where each dataset contains the profile of gene expression, protein and chromatin accessibility, respectively for Y, X and A.

The *α* parameter transforms the samples into the same reduced dimensional plane along the sample space. Furthermore, the *β* parameter transforms the features across the datasets into the same space. The matrix *αX* are the **encoded features** where the sample space has been encoded leaving only the features in the original space. The matrix *Xβ* are the **encoded samples** where the feature space has been encoded leaving only the samples in the original space.

### 2.1 Further details of the Generative Encoder

Inspection of the parameters and how they are linked can be taken a further step. By expressing *Y, X*, and *A* as a function of the parameters, it is possible to extract the core common element shared between all datasets called the code. For example, let the code be *Z* of reduced dimension *k* by *c*.

For *Y* the decoded estimate is *Ŷ*,

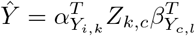

Likewise, *X* can be recovered by the decoded estimate 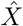

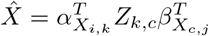

Finally, *A* can be recovered by the decoded estimate *Â*

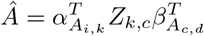

These expressions using the code *Z* can recover the original datasets, with the key important concept that the latent space shared amongst the datasets are given as *Z*.

Furthermore, it can be limiting to consider the datasets as matrices - they can be of any size, including three dimensional objects or tensors.

### 2.2 Properties of the Generative Encoder

#### Functions and matrix properties

The encoding can be considered a learned set of functions which operate on the data modalities. Let *α* and *β* represent *g*(.) and *f* (.) respectively, such that going from left to right:

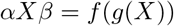

Note it is important to learn the optimisation computationally such that

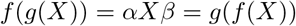

Which is the property of associativity.

#### Samples can be encoded

Sample sizes can be reduced and thus save computational resources when running prediction or nearest-neighbour algorithms. For example, by first reducing the large sample size (several million data points) to a smaller encoded set (several thousand), an algorithm can be run on the smaller set and then decoded back into the original space.

#### Data can be recovered and imputed

Given all data points can be fully expressed by the model, then iterating over updates of the model while simultaneously updating any missing data points can recover the structure of the data and impute missing points. This is similar to the concept of a bootstrap, yet for prediction.

#### Transformations can be transferred

If of similar sample ID the sample parameter *α* can be transferred, if of similar feature label the feature parameter *β* can be transferred. For this reason, a transform learned to reduced the dimensions from a set of genes can be trained on one dataset, and transfered to the second dataset, provided the gene list is identical.

#### Parameters can be joined

If multiple datasets share the same sample ID, the sample parameter can be shared. Otherwise, if multiple datasets have overlapping feature labels, the feature parameter can be shared. This provides an extra level of interpretability.

### 2.3 Coordinate descent updates for learning parameters

The coordinate descent update to estimate the parameters, iterates through multiple steps outlined by Algorithm 1 until convergence. The full step loops over all datasets, and is run in a while loop within gcode.

#### Algorithm 1 Generative Encoding via Generalised Canonical Procrustes

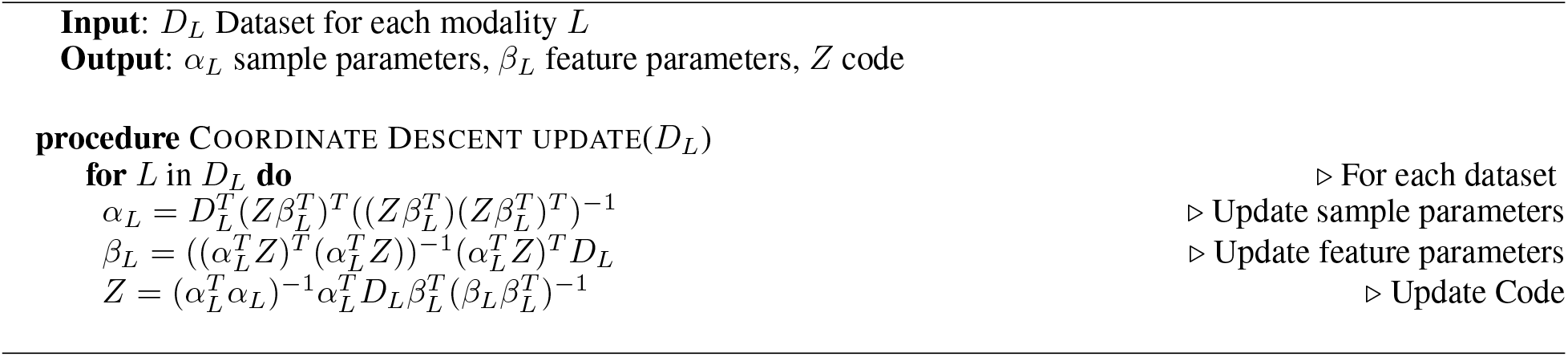

### 2.4 Why learn in a subspace?

Learning in a reduced subspace enables more computational efficiency as the parameters are learned in the smaller reduced dimension. Furthermore, when the dimensions of both sample and features are encoded down into a reduced dimension, it is possible to learn an algorithm on the encoded data, and then project the learned information back up into the original space via a decoding - with results comparable to when the algorithm is run on the full samples.

## 3 Numerical Results

The results contains two parts - a run on single cell data, and, an integration of histology images with RNA-seq data:

### 3.1 Clustering and Trajectory of RNA from single cells of adult mouse hippocampus

Generative encoding summarises information by encoding the sample space into a reduced dimension (in addition to encoding the feature space). Doing so reduces the samples to a smaller dimension, capturing and then representing only the most important structures in the data. This reduces variability and concentrates points to identifiable clusters in visualisation dimension reduction tools such as UMAP or t-SNE.

Here single cell data from an adult mouse hippocampus has had transcriptome measured at a nuclei level [9]. It is expected for both visualisations: 1 - with, and 2 - without gcode; to represent the anatomical structure of the adult mouse hippocampus.

In Figure 2, cell types are more visible - which is expected given gcode encodes the sample points to refine structure. Clusters of points exhibit the finer structures of cell types: clustering well with the upper anatomical cell labels, and still exhibiting neighbouring proximity to similar cell types in the lower level anatomical annotations.

**Figure 1:**
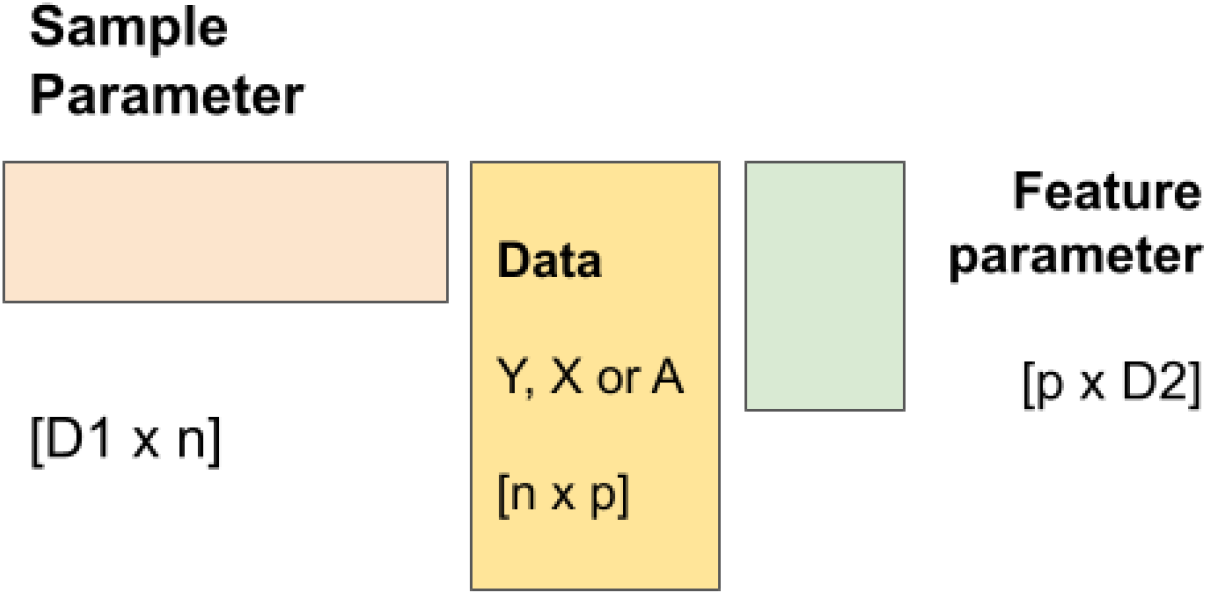
Encoding the dataset using two sets of parameters - an encoding of the sample space and an encoding of the feature space. Multiple sets of data can be aligned to the same space, provided there is a proportional relationship between the two encoding spaces, for example, an equality

**Figure 2:**
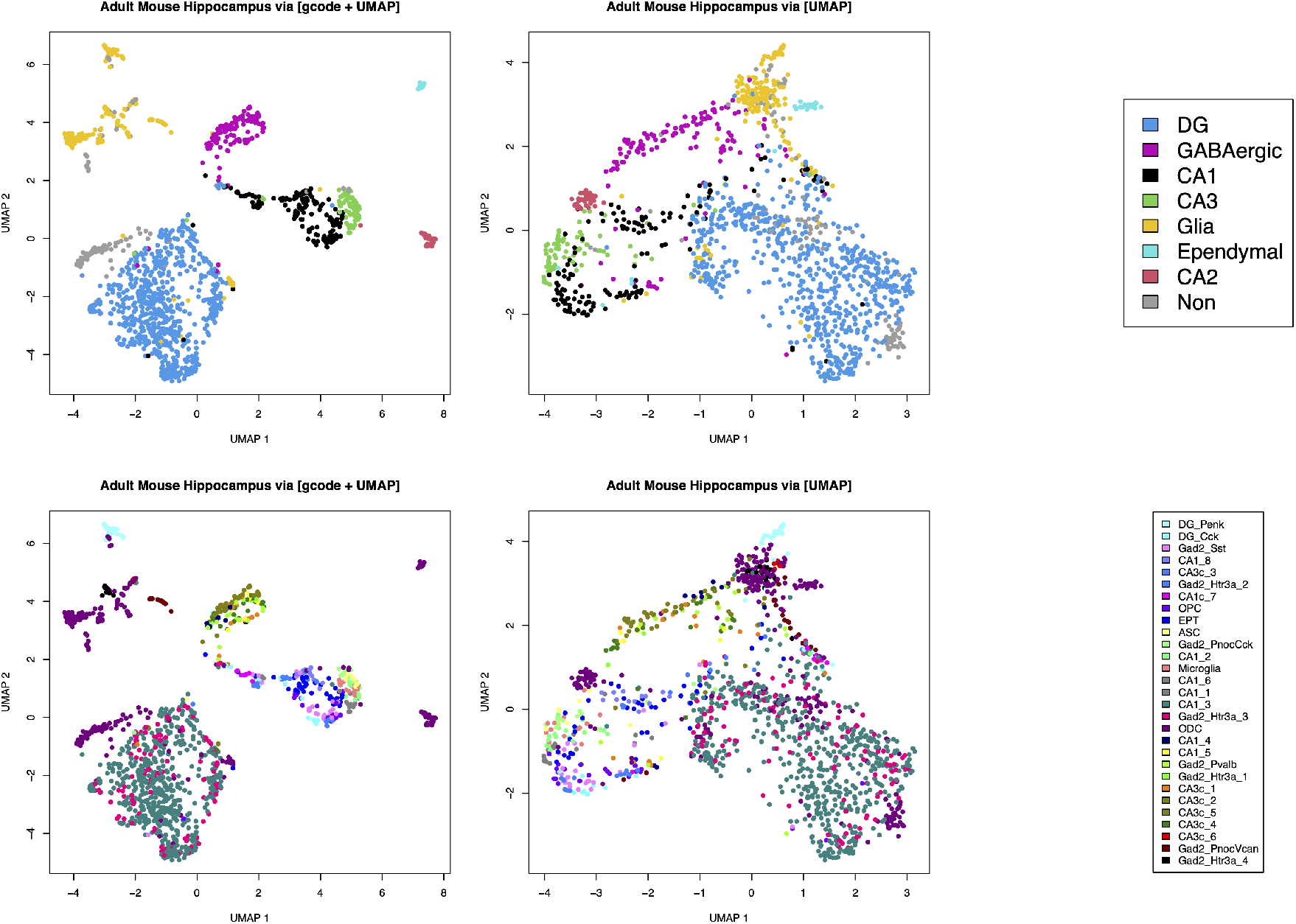
UMAP plots comparing effects of gcode (left) and without (right). Cell type labels are closer neighbours and more visible as clusters for both upper and lower cell annotations.

**Figure 3:**
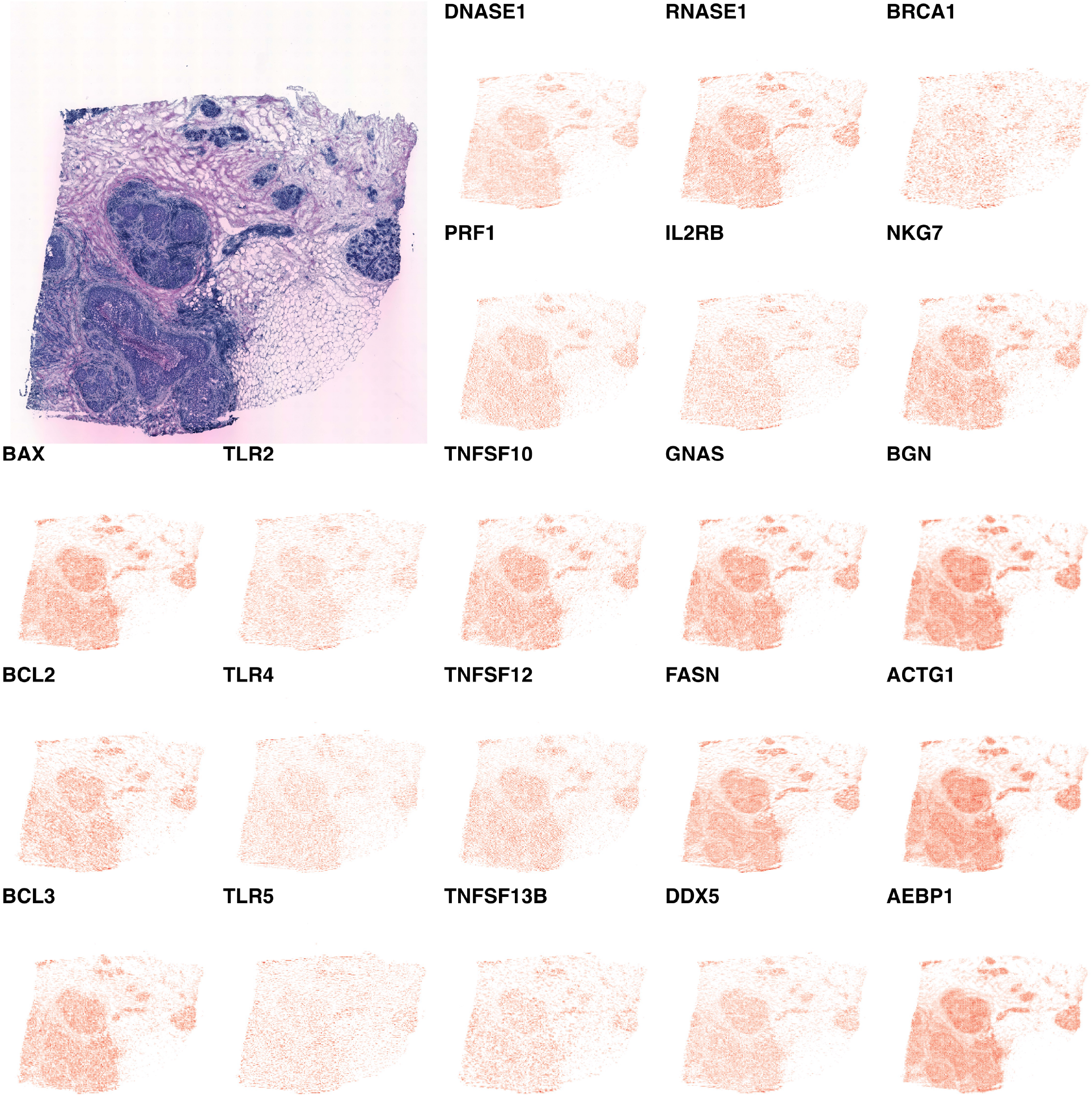
Tumourigenic breast tissue with histology measured using Spatial Transcriptomics accompanied by expression heatmaps of genes. Importantly, genes have been transformed to a standard normal such that relative expression level between genes are at different scales.

### 3.2 Transforming histology to gene expression

Tumourous Breast tissue was taken from He et al, where histological samples were measured with gene expression using Spatial Transcriptomics [10]. Generative Encoding was used to find a transformation between the spatial gene expression and the tissue histology of the Spatial Transcriptomics dataset.

In Figure 4, histology is transformed to gene expression. This idea of finding gene expression values from histology images using spatial cell structure and morphology from Spatial Transcriptomic data is also shared with the work in stNet and XFuse [10] [11].

**Figure 4:**
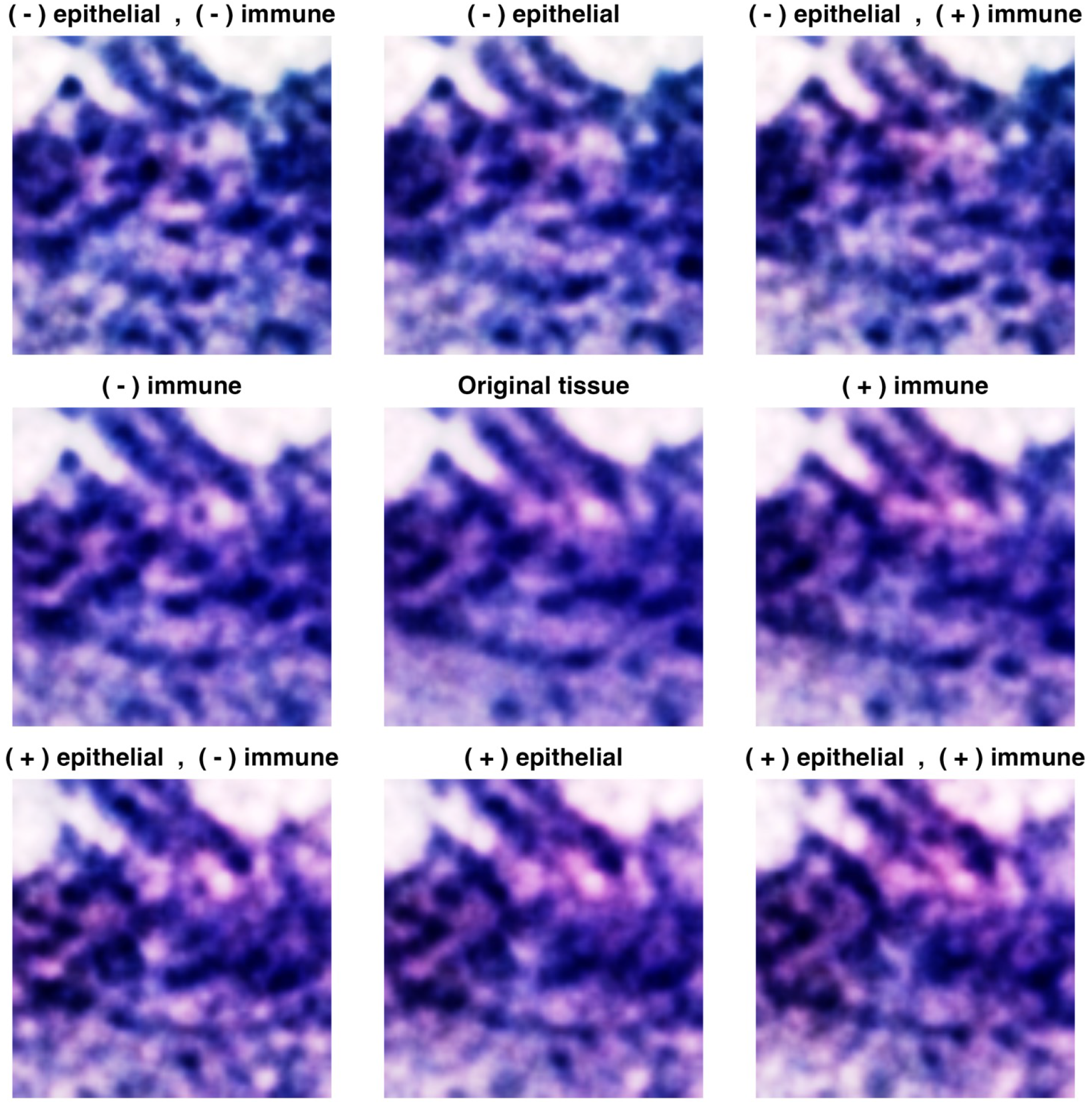
A single image of a H&E stained histology tissue capturing cells in dark blue globules, and extracellular matrix in shades of purple. The center plot shows the original histology image. The vertical and horizontal axis shows the histology image altered by perturbing the expected gene expression of the image (histology transformed to gene expression vector, perturbed, and then transformed back to histology image). From top to bottom shows increasing expression of epithelial gene markers, whereas left to right shows increasing expression of immune gene markers.

To begin, let *E* represent the spatial transcript data, and, *H* to indicate the histology pixel matrix information. Now, given the Generative Encoder relation:

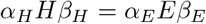

It can be considered that the sample space (represented by *α* is identical between spots of histology pixels, such that it no longer contains relevant information and can be ignored:

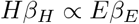

From here, using this relation it is possible to write *H* as the full expression:

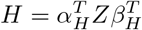

Putting the full expression into the relation for *H*, and the equivalent full expression for *E* gives:

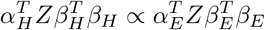

Following, the pixel and gene effects are removed as *β*^*T*^ *β* is orthogonal when the model is initialised via truncated SVD (irlba), and the model is designed to maintain the orthogonality of the parameters:

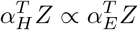

Finally, 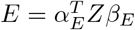 can be regenerated on the right hand side by multiplying with *β*_*E*_:

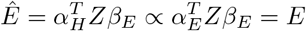

This gives the relation and the eventual transformation of the histology *H* to the expected expression of the histology *Ê*, which is directly proportional to *E*. Through 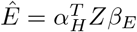, the samples of *Ê* are described by *α*_*H*_, the histology sample parameter.

The large histology image slide is tiled into a space interval of 100 pixels, with each tile of 100 × 100 RGB pixels. These are then transformed to gene expression vectors, whereby heatmaps can be generated. The heatmap is plotted such that the expression is scaled first by a z transform along the genes to a standard normal distribution:

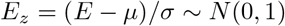

Where *E* indicates gene expression data and *E*_*z*_ indicates the scaled gene expression.

Following on, a relative transform *t* by the minimum and maximum such that the lowest expression value is closest to -1 and the highest expression value is closest to +1, although, not necessarily equal to -1 or +1. By first transforming to a standard normal, it allows different genes to be compared relative to each other according to their distributions:

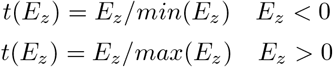

Notice in Figure 4 that the detailed patterns exhibited by gene heatmaps have a diverse, variable, and highly detailed structure. When comparing genes that are expressed in extracellular matrix - BGN, ACTG1 or AEBP1, notice the patterns are very smooth near dense regions of cells. The BCL and TLR gene families show a rough patterning, with small striped lines going mostly in a single direction. Other genes representing Natural Killer cells, PRF1, IL2RB1, and NKG7 are sparser and reveal spotted regions of activity with other regions of inactivity.

### 3.3 In-silico perturbation of histology tissue with gene expression mappings

Histology images can be transformed to gene expression vectors for each image, followed by a perturbation to alter the gene expression level relative to each gene by a factor of the gene expression standard deviation representing the density required for approximately 25% to cover a two-tailed standard normal distribution.

A transformation similar to the histology to gene expression projection is used from the previous section.

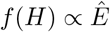

For any given gene *s* in a gene set *G*, the perturbation 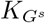 is calculated via *ϕ*^*−*1^ which is the inverse cumulative distribution function of a normal distribution with probability *p* = 25%.

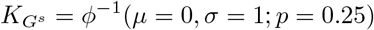

The final perturbation on genes *G*^*s*^ is a addition or subtraction by 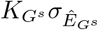, where *σ* is the standard deviation,

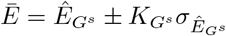

Thereafter, the perturbed expression *Ē* is transformed back into a histology image. This step follows the same processes in the first experiment when converting histology to gene expression although with the inverse of the transform.

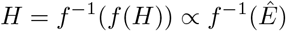

Notice in Figure 4, with increasing epithelial gene expression, the tissue is increasingly marked for a pink eosin stain. Epithelial and immune marker genes are shown in the Supplementary.

## 4 Related Work

### 4.1 Similarities to Singular Value Decomposition

When the model is expressed as *αY β* it is similar to a Procrustes Analysis. However, the detailed model outlined in further detail (Section 2.1), *Y* = *α*^*T*^ *Zβ*^*T*^ shows an expression depicting Singular Value Decomposition.

### 4.2 Similarities to Procrustes

In order to align two sets of shapes or points within a dataset, the method centers and scales the points within each set, and then takes a reference set upon which the other set is rotated and fit to. The expression for the centering and rescaling with rotation is : *Y*_*i,l*_ = *k*_*k,i*_*X*_*i,j*_*Q*_*j,c*_ [4].

### 4.3 Similarities to Canonical Correlation Analysis

The aim is to find a set of component vectors for each of the reference and the non-reference dataset which maximise the correlation. The generative encoder borrows ideas, by finding sets of parameters for each dataset that maximises the similarity of the structures of the encoded datasets.

### 4.4 Similarities to Neural Networks and Autoencoders

Deep Learning models are optimisation methods that use a non-linear composition of weights to minimise a loss function [2]. The style of using many weights that scales with both features and samples to analyse images is capable with both Deep Learning and Generative Encoding.

## 5 Conclusion

Encoding datasets into a shared subspace enables the simultaneous analysis of all datasets. The analysis is done through learned data transformations that are parameters which enable a functional mapping between modalities.

The properties of the Generative Encoder are shown in the summarising ability to concentrate signal and reduce spurious noise. The data transformation from Generative Encoding is evaluated by mapping between histology and gene expression. An example of the mapping is the transform from images of a histology tissue specimen into a gene expression heatmap at a detailed level. Another example is the perturbation of a given histology image, by perturbing the expected gene expression of any given tissue image.

The study shows the capacity of Generative Encoding to be an interpretable analysis tool for biologists and in other disciplines where there is a need to make use of many datasets of different modalities.

## 6 Code and Data

The GitHub version of the gcode package: https://github.com/AskExplain/gcode/

The GitHub scripts used to output figures and analyses: https://github.com/AskExplain/gcode_analysis

More figures on the in-silico perturbation and transforamtion of gene expression to histology: https://doi.org/10.5281/zenodo.5851702

The GitHub verison of R for gcproc (previous version of gcode): https://github.com/AskExplain/gcproc/

A preliminary GitHub version of Python for gcproc (previous version of gcode): https://github.com/thisismygitrepo/gcprocpy

A dashboard detailing ideas on the role of Generative Encoding with Spatial Transcriptomics: https://board.askexplain.com/genecode

## 7 Acknowledgements

David Banh would like to acknowledge:

1. Ryan Deslandes (University of Queensland, Australia) for assistance with software development of the genecode dashboard: https://board.askexplain.com/genecode. This idea was presented in the NanoString Hackathon earning third place and bonus prize for best images (first and second place went to the teams from Dr Omer Bayraktar’s Lab from the Wellcome Sanger Institute, one of which presented cell2location).
2. Cameron Gordon and Olivia Ou (University of Adelaide, Australia and University of Queensland, Australia) for consistent support and advice throughout the project.
3. Alex Alsaffar (University of Queensland, Australia) for relevant discussions on the model of Generalised Canonical Procrustes and Generative Encoding as well as work on a preliminary Python version of gcproc (a prior version of gcode): https://github.com/thisismygitrepo/gcprocpy.
4. Dr Quan Nguyen (University of Queensland, Australia) for a brief discussion on the model of corevec (a prior version of gcproc, which is an earlier version of gcode) regarding imputation and alignment in early 2021 https://github.com/AskExplain/corevec

## 8 Supplementary

Epithelial and Immune gene sets that are altered in the in-silico perturbation of histology images are:

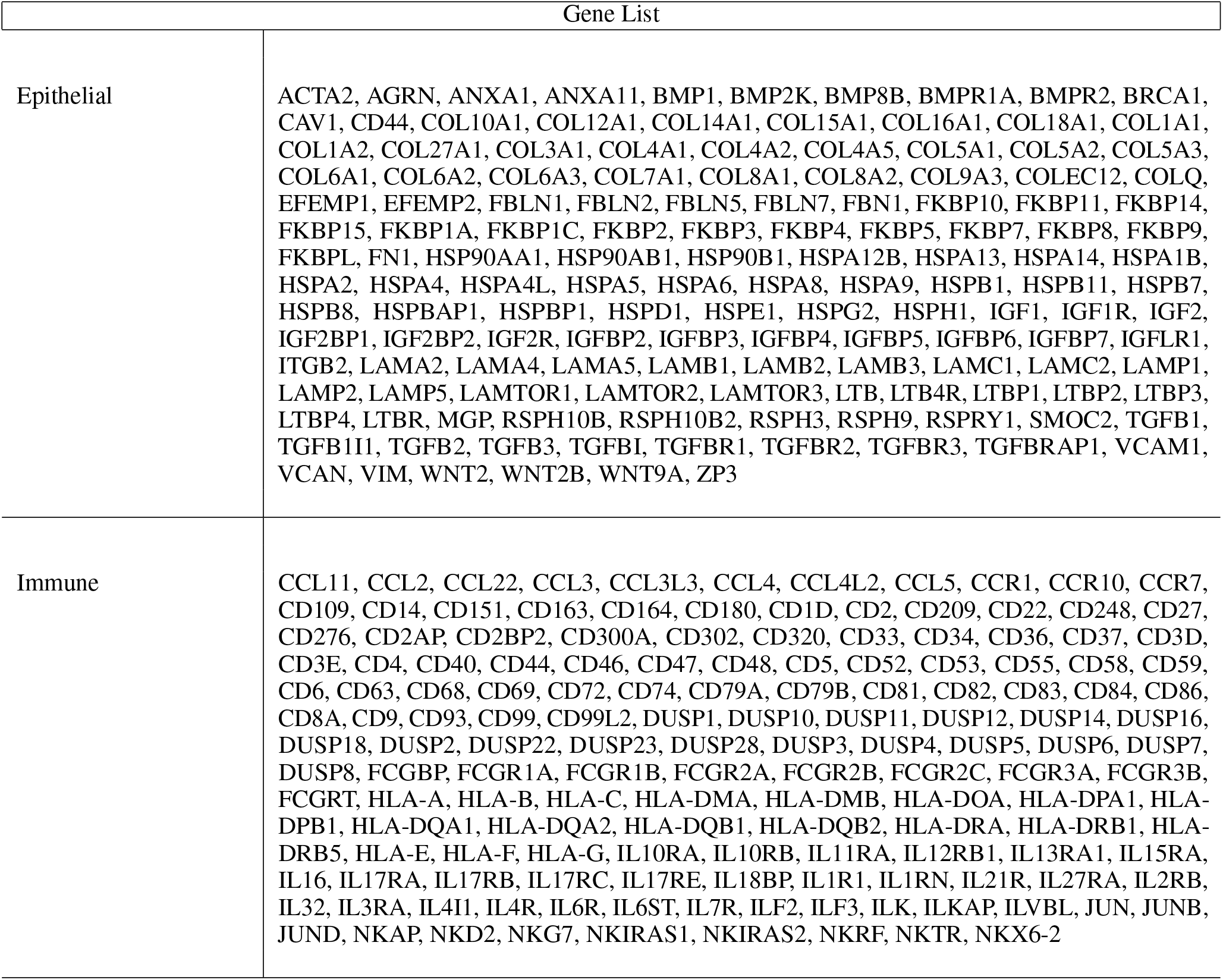

